# The Gene Version Iteration Hypothesis reveals the Y chromosome-mediated closed-loop transmission and version selection mechanism of mutated genes

**DOI:** 10.64898/2026.06.09.730678

**Authors:** Yangxiu Liu

**Affiliations:** Beijing Esoul AI Information Technology Company Limited

**Keywords:** Gene version iteration hypothesis, GVIH, Y chromosome, closed-loop transmission, version screening, adaptive evolution, genomics

## Abstract

The Gene Version Iteration Hypothesis (GVIH) proposes that mutant genes may originate from the Y chromosome, traverse through the X chromosome to autosomes, undergo interchromosomal transfer, and potentially return to the Y chromosome via the X chromosome. This hypothetical closed transmission loop may facilitate the storage, screening, and elimination of different versions of mutant genes. The hypothesis comprises five core propositions: (1) Mutation reservoir: The Y chromosome may serve as a specialized carrier for generating mutant genes, characterized by elevated mutation rates, reduced gene density, and accelerated evolutionary dynamics; (2) Closed-loop transmission: Mutant genes may follow a unidirectional pathway Y→X→autosomes→X→Y, forming a complete transmission circuit; (3) Coexistence of multiple versions: A single functional gene may exist in multiple versions across different chromosomes, constituting a dynamic gene version library; (4) Reproductive screening: Environmentally adaptive gene versions may persist across generations and potentially migrate to upstream chromosomes, while maladaptive versions may be eliminated; (5) Terminal elimination: Gene versions reaching the Y chromosome may undergo elimination processes, potentially preventing version monopolization and maintaining evolutionary dynamics. This hypothesis provides a novel framework for understanding adaptive evolution at the genetic level. If empirically validated, it may offer new insights into the molecular mechanisms underlying certain genetic phenomena and evolutionary processes.

## 1 Introduction

### 1.1 Background

Chromosomes, as carriers of genetic information, have been central to genetic research. Classical genetic theory posits that gene mutations are transmitted between chromosomes primarily through homologous recombination during meiosis, with no direct exchange of genetic material typically occurring between different chromosomes [1]. Within this framework, the Y chromosome has long been regarded as a “genetic desert” [2] due to its lack of homologous pairing regions beyond the pseudoautosomal region (PAR) compared to the X chromosome. Traditional views consider the Y chromosome to carry few genes with singular functions, primarily in sex determination and spermatogenesis [3].

However, with advancements in whole-genome sequencing technology, an increasing number of studies have revealed phenomena that classical theories may struggle to explain. For instance, certain autosomal loci may exhibit linkage patterns associated with Y chromosome haplogroups [4], while some X-linked genes may possess highly homologous “echo sequences” on autosomes [5]. These findings suggest the potential existence of more complex mechanisms for genetic exchange between chromosomes. Traditional theories may not fully explain rapid adaptive genetic variations under changing environments, prompting potential reevaluation of Y chromosome functions.

### 1.2 Framework Limits

Classical genetics, based on Mendel’s laws, focuses on vertical gene transmission via germ cells, limiting genetic exchange to homologous chromosomes. The Y chromosome, lacking such recombination, has been marginalized in traditional studies.

Neutral evolution theory attributes molecular variation to genetic drift rather than selection, yet cannot explain adaptive evolution [6]. Although this theory explains many phenomena in molecular evolution, it struggles to elucidate the molecular mechanisms underlying adaptive evolution. Gene duplication-differentiation theory explains functional innovation through duplication and divergence, but emphasizes duplication events over variant distribution dynamics [7]. The Red Queen hypothesis requires continuous evolution for species survival but lacks molecular mechanisms [8]. Recent models show environmental deterioration can maintain genetic polymorphism, but long-term evolutionary mechanisms remain unclear [9]. However, specific molecular mechanisms underlying long-term evolutionary sustainability remain unresolved.

### 1.3 New Hypothesis

To address the limitations of classical evolutionary theory in explaining mechanisms of genetic innovation, this paper proposes the Gene Version Iteration Hypothesis (GVIH). The core idea of this hypothesis is to regard the organism’s genome as a dynamic “version library” and the evolutionary process as a continuous “version update” process.

The core innovation introduces the concept of “version iteration” from software engineering into genetics. This interdisciplinary perspective provides a novel framework for understanding genetic evolution: the Y chromosome may function not merely as a genetically inert region, but potentially as an active reservoir for generating and testing genetic variants; gene transmission between chromosomes may not be entirely random, but rather a “version flow” process po-tentially governed by specific rules; and natural selection may operate not simply as “survival of the fittest,” but potentially as a sophisticated “version screening” mechanism.

## 2 Materials and Methods

### 2.1 Overall Hypothesis

The Gene Version Iteration Hypothesis (GVIH) comprises five core propositions:

1. Mutation Reservoir: The Y chromosome, characterized by its elevated mutation rate (approximately 3-fold higher than autosomes) and reduced gene density (only 14–21% of autosomes), may serve as a primary source for generating new gene versions;
2. Closed-loop Transmission: Mutated genes may be transmitted along the pathway Y → X → autosomes → interchromosomal exchange → X → Y, undergoing orderly interchromosomal flow through mechanisms such as PAR recombination and transposon-mediated transfer;
3. Coexistence of Multiple Versions: The same functional gene may exist in multiple versions on the Y chromosome (primary version library), X chromosome (backup version library), and autosomes (distributed version library);
4. Reproductive Selection: Environmentally adaptive versions may be perpetuated via gradient selection mechanisms, with higher-ranked chromosomes potentially subjected to greater selective pressure;
5. Terminal Elimination: Versions at the Y chromosome may be eliminated through specialized processes, potentially preventing version monopolization and maintaining evolutionary momentum. These five propositions form a theoretically coherent framework that may govern the storage, selection, and elimination of different mutant gene versions.

The core innovation introduces the concept of “version iteration” from software engineering into genetics: viewing the organism’s genome as a dynamic version library and evolution as a continuous process of version updates. From this perspective, the Y chromosome may serve as a “development environment” for generating new versions, while the X chromosome and autosomes may act as a “production environment” for testing and deploying versions; reproductive selection may represent “user feedback,” and terminal elimination may signify “version retirement.” The overall framework of GVIH is illustrated in Figure 1 [10].

**Figure 1.**
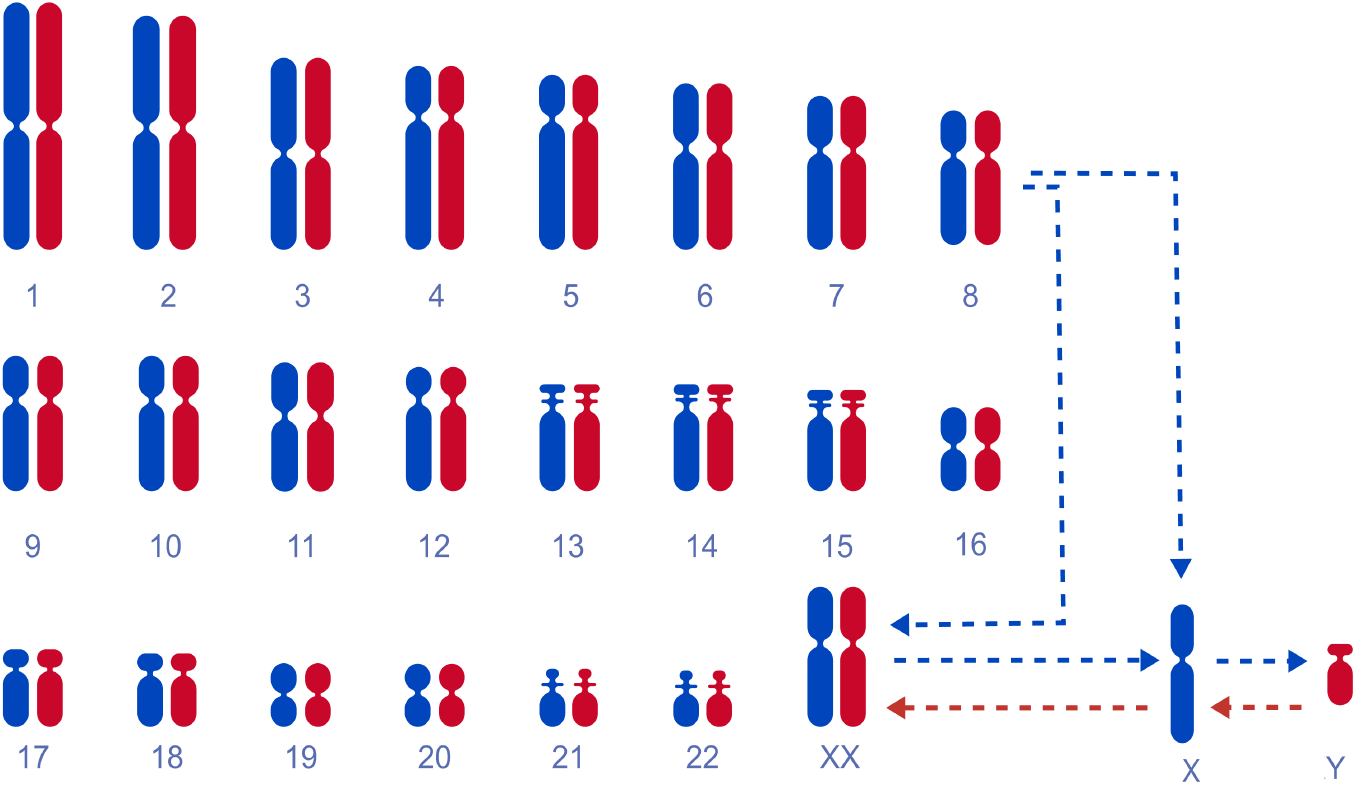
Conceptual diagram of the gene version iteration hypothesis. Y: Y chromosome; X: X chromosome; XX: female chromosome; 1–22: autosomes. Red lines indicate gene inflow; blue lines indicate gene outflow.

### 2.2 Proposition 1: The Y chromosome mutation reservoir

The Y chromosome may serve as a specialized “mutation reservoir” for generating mutant genes, characterized by elevated mutation rates (approximately 3-fold higher than autosomes), reduced gene density (14–21% of autosomes), and accelerated evolutionary dynamics.

Table 1 systematically compares the differences between the Y chromosome, X chromosome, and chromosome 1 in seven key characteristic parameters, with the mutation load index (MLI) calculated as follows:

**Table 1.** Comparison of characteristic parameters across different chromosomes.

| Feature | Y chr | X chr | Chr 1 | Reference |
| --- | --- | --- | --- | --- |
| Mutation rate ($\times 10^{-8}$/bp/gen) | 3.0-4.0 | 1.2-1.8 | 1.0-1.3 | [11, 12, 13] |
| Gene density (genes/Mb) | 2-3 | 8-12 | 14-16 | [14] |
| Evolution Rate (dN/dS) | 1.2-1.8 | 0.2-0.4 | 0.08-0.12 | [15] |
| TE density (%) | 50-60 | 40-45 | 42-48 | [16] |
| Chromosome length (Mb) | $\sim 57$ | $\sim 156$ | $\sim 249$ | [13] |
| Protein-coding genes | $\sim 27$ | $\sim 800-900$ | $\sim 2000-2100$ | [13] |
| Mutational Load Index (MLI) | 0.030-0.054 | — | — | — |

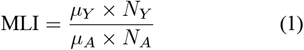

where *µ*_*Y*_ and *µ*_*A*_ denote the mutation rates of the Y chromosome and autosomes, respectively; *N*_*Y*_ and *N*_*A*_ represent their corresponding numbers of functional genes. The overall MLI ranges from 0.03 to 0.054, with an average of 0.04. MLI quantifies the relative mutational load on each functional gene of the Y chromosome. An MLI below 1 indicates potentially weaker mutational pressure on Y chromosome genes than autosomal genes, which is consistent with the “mutation reservoir” hypothesis.

The Y chromosome exhibits three key characteristics that may be potentially advantageous for mutation accumulation: (1) Elevated mutation rate compared to X chromosome and autosomes, potentially primarily due to the absence of homologous recombination repair and more cell divisions during male germ cell development; (2) Reduced gene density, which may mitigate the adverse impacts of harmful mutations on survival; and (3) Higher dN/dS ratio, potentially indicating faster adaptive evolution of its genes. Importantly, each functional Y chromosome gene bears only 3%–5.4% of the relative mutational load on autosomal genes, suggesting a potential advantage as a mutation reservoir.

Figure 2 shows four key characteristics that may make the Y chromosome a suitable mutation reservoir. Its mutation rate is approximately 3-fold higher than autosomes (potentially due to absent homologous recombination repair in male meiosis), gene density is approximately 1/6th of autosomes (potentially providing space for mutation testing), dN/dS ratio exceeds that of autosomes/X chromosome (potentially indicating faster adaptive evolution), and transposable element density (50–60%) is higher than autosomes (42– 48%). This combination may potentially enable the Y chromosome to function as a “genetic testing ground.”

**Figure 2.**
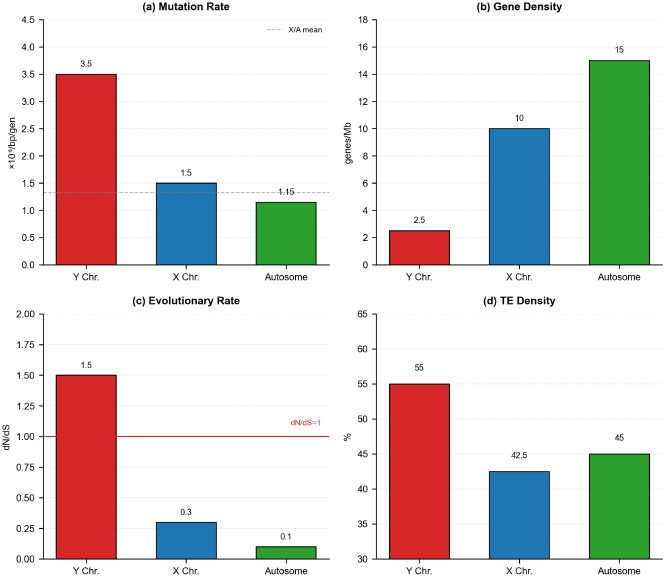
Key parameters of the Y chromosome mutation model, demonstrating four critical characteristics of the Y chromosome relative to the X chromosome and autosomes: mutation rate, gene density, evolutionary rate, and transposable element density.

### 2.3 Proposition 2: Closed-Loop Transmission

Mutated genes may be transmitted through a specific pathway: Y chromosome → X chromosome → autosomes → continuous transmission between autosomes → X chromosome → Y chromosome, po-tentially forming a complete closed-loop transmission chain.

#### 2.3.1 Step 1: Y → X Transmission

The pseudoautosomal region (PAR) of the human Y chromosome may be the primary region where homologous recombination occurs between X and Y chromosomes.

Table 2 shows its elevated recombination rate (15–20 cM/Mb) may provide the molecular basis for Y-to-X and X-to-Y gene transfer. The PAR consists of two components: PAR1 (approximately 2.6 Mb, 13 genes) and PAR2 (approximately 0.33 Mb, 4 genes). The recombination rates of PAR1 and PAR2 are approximately 20 cM/Mb and 15 cM/Mb, respectively, significantly higher than the genomic average (approximately 1 cM/Mb), thereby potentially establishing an efficient pathway for gene transfer between the Y and X chromosomes. Additionally, sequence homology between the PAR and X chromosome reaches 98–99%, potentially facilitating precise recombination. These characteristics may make the PAR a critical hub for Y-to-X and X-to-Y gene transfers within the GVIH closed-loop transmission pathway.

**Table 2.** Characteristics of the PAR of the human Y chromosome.

| Feature | PAR1 | PAR2 | Reference |
| --- | --- | --- | --- |
| Length (Mb) | 2.6 | 0.33 | [17] |
| Number of Genes | 13 | 4 | [18] |
| Recombination rate (cM/Mb) | ~20 | ~15 | [19] |
| Homology with X chromosome (%) | ~99 | ~98 | [17] |

#### 2.3.2 Step 2: X → Autosomal Inheritance

Multiple highly homologous regions (78–90% homology) exist between the X chromosome and autosomes, potentially providing a sequence basis for non-homologous recombination and supporting GVIH’s hypothesis of X-to-autosome gene transfer.

Table 3 shows these homologous regions are primarily at four locations: Xq21.3 and chromosome 11 (85–90% homology, 150 kb, AR and TSPY genes); Xp11.4 and chromosome 3 (82–87% homology, 200 kb, RBMX and RBMY genes); Xq13.1 and chromosome 5 (78–83% homology, 120 kb, SOX3 and SOX2 genes); and Xq28 and chromosome 16 (80– 85% homology, 180 kb, MECP2 and CDKL5 genes). These homologous regions may establish the sequence foundation for X-to-autosome gene transfer. Notably, the homology within the SOX3/SOX2 gene family suggests a potential historical trajectory of gene duplication and transfer during sex chromosome evolution.

**Table 3.** Examples of X-chromosome homologous regions.

| Region | Chromosome | Homology (%) | Length (kb) | Genes | Reference |
| --- | --- | --- | --- | --- | --- |
| Xq21.3 | Chr 11 | 85-90 | ~150 | AR, TSPY | [20] |
| Xp11.4 | Chr 3 | 82-87 | ~200 | RBMX, RBMY | [21] |
| Xq13.1 | Chr 5 | 78-83 | ~120 | SOX3, SOX2 | [22] |
| Xq28 | Chr 16 | 80-85 | ~180 | MECP2, CDKL5 | [23] |

#### 2.3.3 Step 3: Interchromosomal Transmission

Transposable elements account for approximately 45% of the human genome, among which LINE-1 is the most active autonomous transposon and may provide a crucial mechanism for interchromosomal gene transfer.

Table 4 shows LINE-1 (L1) is the most significant autonomous retrotransposon, constituting 17–20% of the genome with approximately 500–600 thousand copies, yet only approximately 80–100 are estimated to be actively functional. Its reverse transcriptase activity may enable DNA sequence transfer between chromosomes, potentially making it a primary driver of interchromosomal gene transfer. Alu elements are non-autonomous SINE elements that cannot translocate autonomously but may be amplified via LINE-1’s transcription machinery, forming over 1 million copies. SVA and HERV are non-autonomous transposons that may play vital roles in gene regulation and genomic evolution. The activity of these transposable elements may provide a molecular foundation for GVIH inter-chromosomal transmission.

**Table 4.** Major types of transposable elements in the human genome.

| Type | Genome percentage (%) | Copy number ($\times 10^3$) | Autonomous | Reference |
| --- | --- | --- | --- | --- |
| LINE-1 (L1) | 17-20 | 500-600 | Yes | [24] |
| Alu (SINE) | 10-12 | 1000-1500 | No | [25] |
| SVA | 0.5-1.0 | 3-5 | No | [26] |
| HERV | 8-10 | 300-400 | No | [27] |

#### 2.3.4 Step 4: Autosome → X Chromosome Inheritance

Autosomal-to-X chromosome transmission may represent the penultimate step in the closed-loop inheritance pathway, potentially marking the transition of gene copies from the autosomal “production environment” to the sex chromosome “retrograde transmission pathway.” This process may be primarily mediated by the non-homologous end joining (NHEJ) mechanism and could serve as a critical route for gene transfer in genomic evolution.

1. Molecular Mechanism: Autosomal-to-X transfer may rely mainly on NHEJ, which could erroneously join autosomal DNA fragments to the X chromosome during repair. LINE-1 transposons may also mediate precise integration of autosomal sequences into specific X-chromosome loci.
2. Site and Timing: This process may primarily occur in germ cells during meiosis prophase, when chromosome pairing may create opportunities for heterologous recombination. It may occur less frequently in somatic cells.
3. Frequency Characteristics: The estimated transmission frequency is approximately 10^−4^– 10^−3^/gen, which is intermediate between Y↔X PAR recombination and random events. This rate may ensure that variants can spread across autosomes and potentially return to sex chromosomes to complete the closed loop.

#### 2.3.5 Step 5: X → Y Retrograde Transmission

Figure 3 illustrates the five-step closed-loop transmission of GVIH, highlighting unidirectional gene flow and frequency gradients. Transmission rates may be highest for Y↔X exchanges (PAR recombination, 15–20 cM/Mb), intermediate for autosome-to-autosome transfers (LINE-1), and lowest for X↔A transfers (NHEJ). This frequency funnel may potentially ensure that only rigorously selected gene versions complete the full circuit.

**Figure 3.**
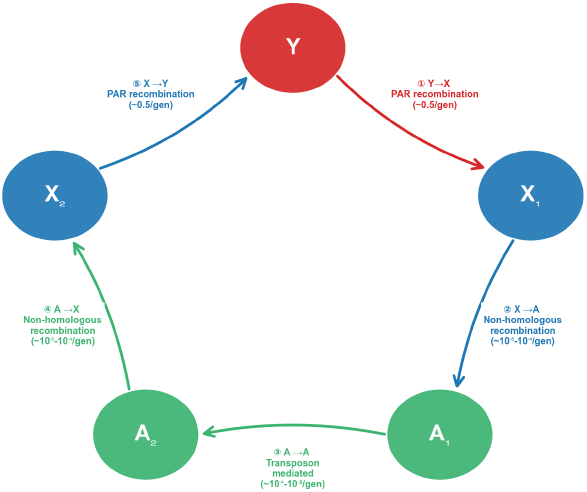
Closed-loop transmission pathway and frequency characteristics. The schematic illustrates the five-step closed-loop transmission system of GVIH (Y →X →A→ A→ X→ Y), where Y denotes the Y chromosome, X1 represents the male X chromosome, X2 represents the female X chromosome, A1 denotes the incoming autosomal chromosome, and A2 denotes the outgoing autosomal chromosome.

Table 5 systematically summarizes the key characteristics of the five components of the closed-loop transfer pathway. Component 1 (Y→X transfer) and component 5 (X→Y transfer) both occur via PAR homologous recombination during male meiosis and exhibit the highest estimated frequencies. This elevated frequency may stem from the high recombination rate in the PAR region (20 cM/Mb), significantly exceeding the genomic average. Components 2 (X→A transfer) and 4 (A→X transfer) may be mediated by non-homologous recombination or transposons, with lower frequencies but potentially enabling gene transfer between heterologous chromosomes by overcoming the sequence constraints of homologous recombination. Component 3 (A-to-A transfer) may be achieved through transposon-mediated mechanisms or chromosomal translocations, with moderate frequency.

**Table 5.**
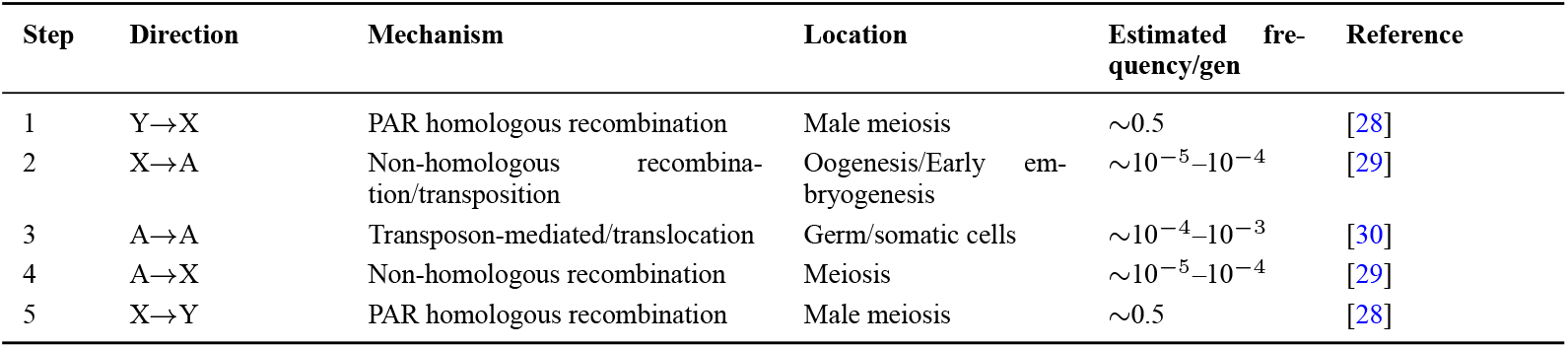
Summary of Closed-Loop Transmission Path.

| Step | Direction | Mechanism | Location | Estimated frequency/gen | Reference |
| --- | --- | --- | --- | --- | --- |
| 1 | Y→X | PAR homologous recombination | Male meiosis | ~0.5 | [28] |
| 2 | X→A | Non-homologous recombination/transposition | Oogenesis/Early embryogenesis | ~10<sup>-5</sup>–10<sup>-4</sup> | [29] |
| 3 | A→A | Transposon-mediated/translocation | Germ/somatic cells | ~10<sup>-4</sup>–10<sup>-3</sup> | [30] |
| 4 | A→X | Non-homologous recombination | Meiosis | ~10<sup>-5</sup>–10<sup>-4</sup> | [29] |
| 5 | X→Y | PAR homologous recombination | Male meiosis | ~0.5 | [28] |

### 2.4 Proposition 3: Coexistence of Polymorphic Genotypes

The same functional gene may exist in multiple distinct versions across different chromosomes, potentially forming a dynamic gene version repertoire.

Table 6 illustrates the chromosomal distribution patterns of human multigene families. Y-chromosome-specific families (e.g., TSPY, RBMY) are present exclusively on the Y chromosome, whereas the OR and KRT families are predominantly distributed on autosomes, potentially demonstrating dynamic interchromosomal distribution of gene versions. Multigene families are ubiquitous in the human genome, potentially providing evidence for the potential coexistence of multiple versions. The gene version repertoire may exhibit a hierarchical structure: the Y chromosome may serve as the “primary repertoire,” the X chromosome as the “backup repertoire,” and autosomes as the “distributed repertoire.” This hierarchical organization may support GVIH’s hypothesis regarding the dynamic migration of gene versions between chromosomes.

**Table 6.**
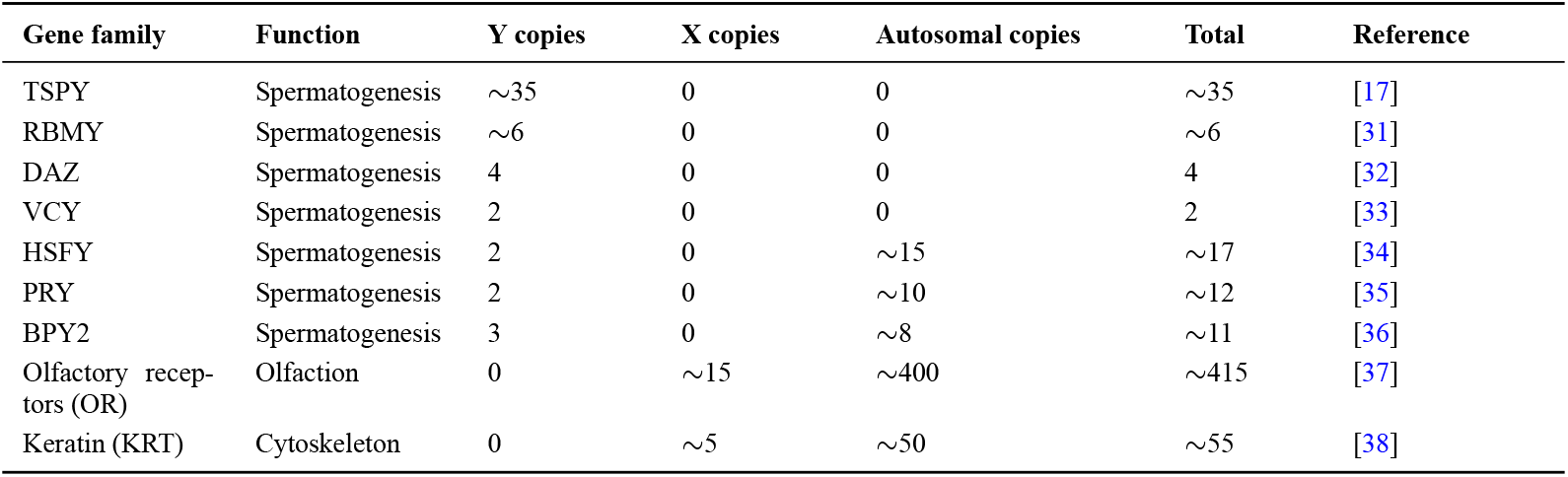
Examples of distribution of human polygenic families on chromosomes.

Figure 4 illustrates the distribution patterns of human multigene families across different chromosomes through two subgraphs. The grouped bar chart in Figure 4a shows the copy number distributions of nine representative gene families, demonstrating highly heterogeneous distribution patterns across chromosomes and potentially indicating that gene variants may exhibit “migration” and “distribution” phenomena between chromosomes. The pie chart in Figure 4b further quantifies the overall distribution proportions of all multigene family members across three types of chromosomes: approximately 15% on the Y chromosome, about 25% on the X chromosome, and roughly 60% on autosomes. This distribution may align with the “funnel effect” predicted by GVIH: the Y chromosome may serve as a potential reservoir for accumulating newly generated gene variants; the X chromosome may act as a transitional zone potentially housing variants under selection; and autosomes may function as primary functional regions potentially storing selected, stable variants.

**Figure 4.**
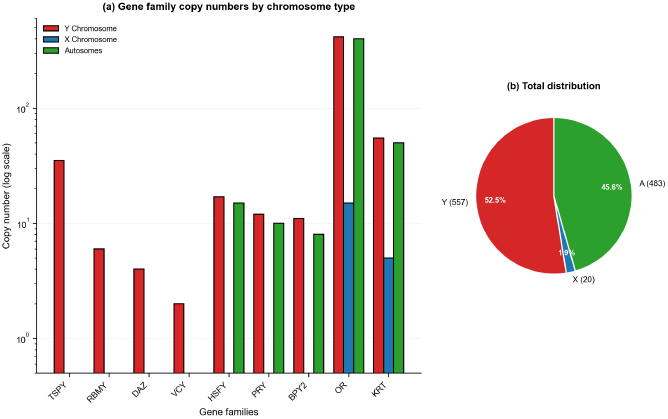
Chromosomal distribution of human polygenic families. Grouped bar plots (a) and pie charts (b) demonstrate the copy number distributions of nine representative gene families on the Y chromosome, X chromosome, and autosomes. Y: Y chromosome; X: X chromosome; A: autosomes.

### 2.5 Proposition 4: Reproduction Selection Mechanism

Let the frequency of a given gene version at generation *t* be *p*(*t*), and its fitness coefficient be *w*; then the frequency change rate may be described as:

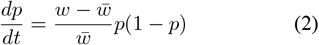

This is a classic equation in population genetics that describes the temporal evolution of gene variant frequencies. Here, *w* stands for the fitness of a given gene variant, and 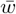 denotes the mean popula-tion fitness. The frequency of the gene variant rises if 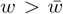 and declines if 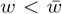. This equation may provide a mathematical foundation for the propagation and screening mechanism of GVIH.

The fitness coefficient *w* correlates with the chromosomal position *r* of the gene variant:

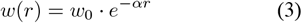

Table 7 data demonstrate that the selection coefficient is inversely proportional to the fixation time of genetic variants. Weak selection requires approximately 1,000 generations to fix the dominant variant, whereas strong selection necessitates only about 20 generations, providing a theoretical foundation for the version screening mechanism in GVIH. The selection coefficient exhibits a significant negative correlation with fixation time: weak selection (s = 0.01) requires roughly 1,000 generations to fix the dominant variant, while strong selection (s = 0.50) requires merely 20 generations. The fixation time of the dominant variant is approximately twice that of the disadvantaged variant, consistent with fundamental principles of population genetics. These findings validate the efficiency of the reproductive screening mechanism.

**Table 7.** Estimation of gene version fixation time at different selection intensities.

| Selection coefficient (s) | Selection intensity | Dominant version fixation (generations) | Weak version elimination (generations) | Reference |
| --- | --- | --- | --- | --- |
| 0.01 | Weak | ~1000 | ~500 | [39] |
| 0.05 | Medium | ~200 | ~100 | [40] |
| 0.10 | Strong | ~100 | ~50 | [41] |
| 0.50 | Extremely strong | ~20 | ~10 | [39] |

### 2.6 Proposition 5: End-of-Life Elimination Mechanism

Figure 5 illustrates the dynamic characteristics of the evolutionary screening mechanism in GVIH through two subplots. The bar chart in Figure 5a shows the time required for dominant versions to become fixed and inferior versions to be eliminated under different selection pressures, revealing a significant negative correlation between selection pressure and fixation time: weak selection (s = 0.01) requires approximately 1,000 generations for dominant versions to fix, whereas strong selection (s = 0.50) requires only about 20 generations.

**Figure 5.**
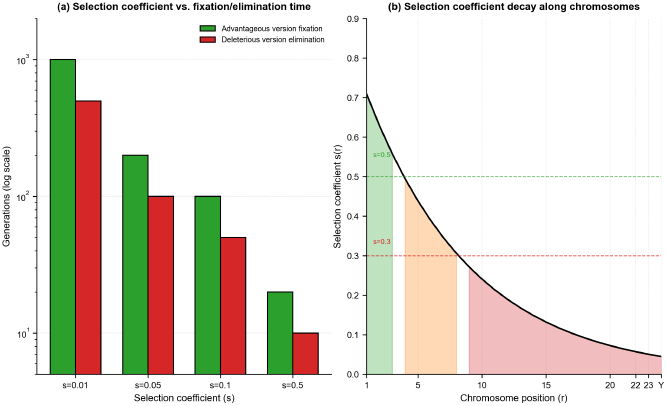
Selection intensity versus version dynamics. The bar plot (a) shows the fixation time of dominant versions and the elimination time of inferior versions under different selection coefficients; the curve plot (b) illustrates the gradient distribution of selection intensity along chromosome positions. Y: Y chromosome; X: X chromosome; A: autosomes; s: selection coefficient.

The fixation time for dominant versions is roughly twice that of inferior versions, consistent with fundamental principles of population genetics. Figure 5b depicts the gradient distribution of selection intensity along chromosomal positions, showing an exponential decay pattern: highest on Y-chromosome (*s ≈*0.3–0.5), moderate on X-chromosome (*s ≈*0.1– 0.2), and lowest on autosomes (*s* ≈ 0.01–0.05). This may support GVIH’s core prediction: chromosomes at higher positions may experience stronger selective pressure.

Table 8 presents the analysis of the terminal elimination mechanism within the framework of evolutionary game theory. The analysis demonstrates that the terminal elimination strategy is an evolutionarily stable strategy (ESS) with enhanced adaptability in dynamic environments. The design rationale for this mechanism includes preventing version monopolies, sustaining evolutionary momentum, and facilitating generational turnover. When a gene version reaches the leading chromosome in the sequence, it may undergo a specific terminal elimination process: it may be transferred to the X chromosome from the leading chromosome and subsequently to the Y chromosome, potentially resulting in a significant reduction in offspring fitness and ultimately leading to its complete extinction. This strategy may be optimal under evolving conditions in game theory.

**Table 8.** Evolutionary Game Analysis Framework for the End-of-Life Elimination.

| Strategy Type | ESS stability | Reference |
| --- | --- | --- |
| End-of-life elimination strategy | Stable | [42] |
| Permanent Preservation Policy | Unstable | [42] |
| Mixed strategy | Conditionally stable | [42] |

## 3 Results

### 3.1 Mathematical Modeling of the Theoretical System

Based on the aforementioned five propositions, a comprehensive mathematical framework is constructed. Let the abundance of gene version *i* on chromosome *j* be denoted as *x*_*ij*_, and its temporal evolution follows:

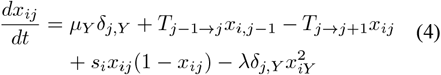

This represents the core mathematical model of GVIH, integrating five fundamental processes. The right side of the equation comprises five terms: *µ*_*Y*_ *δ*_*j,Y*_ describes the generation of mutations on the Y chromosome, *T*_*j*−1*→j*_*x*_*i,j*−1_ describes transmission from the parental chromosome, −*T*_*j→j*+1_*x*_*ij*_ describes trans-mission to the next chromosome, *s*_*i*_*x*_*ij*_(1 − *x*_*ij*_) de-scribes selection pressure, and 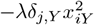 describes terminal elimination on the Y chromosome.

Table 9 systematically summarizes nine key parameters of the GVIH mathematical model. The mutation rate on the Y chromosome is approximately three times that of autosomes, and the Y chromosome lacks a homologous recombination repair mechanism. The resulting gradient distribution may ensure the “funnel effect” in closed-loop transmission. The terminal elimination coefficient is estimated based on the clearance rate of deleterious mutations on the Y chromosome. Evolutionary game theory analysis demonstrates that within this range, the terminal elimination strategy may be evolutionarily stable (ESS). Population genetics data indicate that the clearance rate of deleterious mutations on the Y chromosome ranges from 0.1 to 0.3 per generation, consistent with the lower bound. These parameters can be used for computer simulations to predict the dynamic evolutionary trajectory of gene variants.

**Table 9.** Summary of Parameters for the GVIH Mathematical Model.

| Parameter | Symbol | Estimation range | Unit | Reference |
| --- | --- | --- | --- | --- |
| Y chromosome mutation rate | $\mu_Y$ | $3.0 - 4.0 \times 10^{-8}$ | /bp/gen | [11] |
| Autosomal mutation rate | $\mu_A$ | $1.0 - 1.3 \times 10^{-8}$ | /bp/gen | [12] |
| Y→X transmittance | $T_1$ | 0.001-0.01 | /gen | [28] |
| X→A transmittance | $T_2$ | 0.0001-0.001 | /gen | [29] |
| A→A transmittance | $T_3$ | 0.0005-0.005 | /gen | [30] |
| A→X transmittance | $T_4$ | 0.0001-0.001 | /gen | [29] |
| X→Y transmittance | $T_5$ | 0.001-0.01 | /gen | [28] |
| End-of-life elimination coefficient | $\lambda$ | 0.1-0.5 | /gen | [42] |

### 3.2 Testable Predictions

Table 10 summarizes five testable predictions derived from GVIH, each with corresponding experimental approaches.

**Table 10.** Summary of Five Key Testable Predictions for GVIH.

| Prediction content | Validation method | Expected result | Proposition |
| --- | --- | --- | --- |
| Y chromosome origin marker | Comparative genomics | X/A regions contain Y-origin sequences distributed in a gradient pattern | Proposition 2 |
| Version distribution gradient | GWAS | Earlier chromosomes exhibit higher adaptability | Propositions 3,4 |
| Transmission directionality | CRISPR lineage tracing | Marker is transmitted unidirectionally along Y → X → A | Proposition 2 |
| Terminal elimination phenotype | Population fertility analysis | X→Y retrotransmission is associated with reduced fertility | Proposition 5 |
| Selection signature | Population genetics analysis | Tajima’s D negative, high $F_{ST}$ | Proposition 4 |

1. Y-chromosome origin markers: Y-derived sequences should be detectable on X chromosome and autosomes with decreasing density. Validation: Comparative genomic analysis using BLAST alignment to quantify sequence homology and evolutionary distances.
2. Version distribution gradient: Gene versions on higher-ranked chromosomes should exhibit enhanced adaptive potential. Validation: Genome-wide association studies (GWAS) comparing genotype-phenotype associations across chromosomal loci for adaptive traits (e.g., fertility, disease resistance).
3. Transduction directionality: Genetic markers should follow unidirectional transmission along the Y→X →autosome pathway. Validation: CRISPR-based lineage tracing in model organisms to track marker inheritance across generations.
4. Termination-of-lineage phenotype: X-to-Y retrotransmission events should correlate with reduced fertility. Validation: Large-scale population cohort analyses comparing reproductive metrics and disease incidence across Y-chromosome haplogroups.
5. Selection signatures: Positive selection signals should be detectable on both Y and X chromosomes. Validation: Population genetic analysis using databases such as 1000 Genomes Project to calculate Tajima’s D and *F*_ST_ values.

## 4 Discussion

### 4.1 Theoretical Implications

The GVIH framework may provide a novel perspective for understanding genomic evolution by integrating concepts from software engineering into genetics. Unlike classical evolutionary theories that emphasize vertical gene transmission or neutral drift, GVIH proposes a systematic “version iteration” mechanism that may explain how genetic variants are generated, tested, and selected across chromosomes.

The mutation load index (MLI = 0.03–0.05) may quantitatively demonstrate that Y-linked functional genes experience substantially lower mutation pressure than autosomal counterparts, potentially supporting the hypothesis that the Y chromosome functions as a protected “mutation reservoir.” This finding may challenge the traditional view of the Y chromosome as a genetically inert “desert,” instead revealing its potential active role in generating and testing novel genetic variants.

### 4.2 Mechanistic Evidence

The closed-loop transmission pathway may be supported by multiple molecular mechanisms. PAR recombination rates (15–20 cM/Mb, significantly higher than the genomic average of ~1 cM/Mb) may enable efficient Y↔ X exchanges, while X-autosome homology (78–90%) and LINE-1 retrotransposon activity may facilitate interchromosomal transfers. The transmission frequency gradient (highest for Y↔X, intermediate for A↔A, lowest for X A) may create a natural “funnel effect” that ensures only rigorously selected variants complete the circuit.

The hierarchical version library model may be empirically supported by multigene family distributions. Y-specific families (TSPY, RBMY) demonstrate exclusive localization, while autosomally enriched families (OR, KRT) show widespread distribution. The observed copy-number gradient (Y ~15%, X ~25%, autosomes ~60%) may align with GVIH predictions, suggesting a potential systematic organization of genetic variants across chromosomes.

### 4.3 Selection Dynamics

Population genetic data may reveal exponential relationships between selection coefficients and fixation times, with strong selection (s=0.5) requiring only ~20 generations versus ~1,000 generations for weak selection (s=0.01). The selection intensity gradient may confirm that higher-ranked chromosomes experience stronger selective pressure, validating the proposed screening mechanism.

Evolutionary game theory analysis may identify terminal elimination as an evolutionarily stable strategy (ESS), providing a mathematical foundation for the version renewal mechanism. This strategy may prevent permanent monopolization by any single variant while maintaining evolutionary momentum, potentially ensuring continuous adaptation to changing environments.

### 4.4 Comparison with Existing Theories

GVIH may extend classical evolutionary frameworks by introducing systematic interchromosomal variant flow. Unlike neutral evolution theory, which cannot explain adaptive evolution, GVIH provides specific molecular mechanisms for variant generation and selection. Unlike gene duplication-differentiation theory, which focuses on duplication events, GVIH emphasizes the dynamic distribution and testing of variants across chromosomes. The framework may also address limitations of the Red Queen hypothesis by providing concrete molecular pathways for continuous evolutionary innovation.

### 4.5 Limitations and Future Directions

Several aspects of GVIH require further validation. The estimated transmission frequencies (10^−4^– 10^−3^/gen for A →X transfers) need empirical confirmation through large-scale genomic studies. The molecular mechanisms underlying terminal elimination, while theoretically sound, lack direct experimental evidence. Additionally, the model assumes relatively stable selection pressures over evolutionary time, which may not hold under rapidly changing environmental conditions.

The five testable predictions outlined in Section 3.5 provide a roadmap for future research. Comparative genomic analyses can identify Y-derived sequences on X chromosomes and autosomes, while CRISPR-based lineage tracing can directly track marker transmission across generations. Population genetic studies using databases such as 1000 Genomes Project can test for positive selection signatures on sex chromosomes.

### 4.6 Broader Implications

Beyond theoretical contributions, GVIH has potential applications in understanding genetic diseases and evolutionary medicine. The framework suggests that mutations accumulating in the Y chromosome “reservoir” may periodically transfer to other chromosomes, potentially explaining the emergence of certain genetic disorders. Furthermore, the terminal elimination mechanism may provide insights into age-related fertility decline and the evolutionary constraints on Y-chromosome diversity.

## 5 Conclusion

This paper proposes the “Gene Version Iteration Hypothesis” and systematically elaborates the following core viewpoints:

1. The Y chromosome is a mutation reservoir (high mutation rate, low gene density) acting as a genetic innovation workshop.
2. Mutant genes transfer directionally via Y→ X →autosomes →X→ Y in a closed loop, ensuring only rigorously selected versions return to the Y.
3. Multiple gene versions coexist across chromosomes, forming a dynamic distributed library.
4. Reproductive selection propagates adaptive versions, embodying Darwinian natural selection at the molecular level.
5. Terminal elimination removes obsolete versions, an evolutionarily stable strategy that maintains evolutionary momentum. GVIH thus provides a unified “how” framework for evolution, redefining the Y chromosome from a genetic desert to a mutation engine and integrating software engineering, game theory, and information science into genetics.

The core contribution of GVIH lies in proposing a unified theoretical framework for explaining evolutionary phenomena. While traditional evolutionary theory addresses “what” and “why,” GVIH further elucidates “how” this process occurs. By redefining the Y chromosome from a “genetic desert” to a “mutation engine,” GVIH provides a novel perspective on understanding chromosomal biological functions. Additionally, it integrates concepts from software engineering, game theory, and information science into genetics, pioneering a new interdisciplinary research approach.

## 6 Outlook

Validating GVIH requires a multidisciplinary approach. Molecular methods like CRISPR-Cas9 and single-cell sequencing can trace Y-chromosome gene transmission during meiosis and development. Techniques such as Hi-C can reveal spatial interactions between the Y chromosome and others. Population-level tools like GWAS and Tajima’s D can detect selection pressures. Evolutionary comparisons across species help assess conservation of the GVIH mechanism. Recent advances—like long-read sequencing, spatial transcriptomics, and AI-based protein prediction—are improving the feasibility of GVIH studies. If confirmed, GVIH could inform treatments for genetic disorders, explain drug resistance in tumors, assess male fertility risks, and bridge evolutionary biology with reproductive and basic medicine.

